# Unraveling principles of thermodynamics for genome-scale metabolic networks using graph neural networks

**DOI:** 10.1101/2024.01.15.575679

**Authors:** Wenchao Fan, Chuyun Ding, Dan Huang, Weiyan Zheng, Ziwei Dai

## Abstract

The fundamental laws of thermodynamics determine the feasibility of all natural processes including metabolism. Although several algorithms have been developed to predict the most important thermodynamic parameter, the standard Gibbs free energy, for metabolic reactions and metabolites, their application to genome-scale metabolic networks (GEMs) with thousands of metabolites and reactions is still limited. Here, we develop a graph neural network (GNN)- based model dGbyG for predicting Gibbs energy for metabolites and metabolic reactions which outperforms all existing methods in accuracy, versatility, robustness, and generalization ability. By applying dGbyG to the human GEM, Recon3D, we identify a critical subset of reactions with substantial negative values of the standard Gibbs free energy change, which we name thermodynamic driver reactions. These reactions exhibit distinctive network topological characteristics akin to driver nodes defined in control theory and remarkable heterogeneity in enzyme abundance, implying evolutionary constraints on the selection of thermodynamic parameters of metabolic networks. We also develop a thermodynamics-based flux balance analysis (TFBA) approach to integrate reaction thermodynamics with GEMs to curate these models. Our work not only transcends the augmentation of accessible thermodynamic data to facilitate an enriched understanding of metabolism, but also enables refinement of metabolic reconstructions from a thermodynamic perspective, thereby underscoring the critical role of thermodynamics in the accurate modeling of biological systems.

## Introduction

Thermodynamics is the primal determinant of feasibility of all processes in the nature, including a variety of biological processes on various temporal and spatial scales, from evolution spanning geological epochs^1^ to biochemical reactions^2^ and protein folding and allosteric regulation processes^3, 4^ at the time scale of milliseconds. Recent studies have demonstrated that all aspects of cellular metabolism, including but not limited to the control of metabolic fluxes^5^, the efficiency of metabolic pathways^6^, and the allocation of cellular resource into different metabolic activities^7^, are profoundly affected by the Gibbs free energy, an important thermodynamic parameter of metabolic reactions and metabolites. Moreover, quantitative knowledge about thermodynamic parameters of metabolic reactions is indispensable for accurate reconstruction^8^ and curation^9, 10^ of genome-scale metabolic models (GEMs), which are instrumental for predicting metabolic phenotypes^11–13^, designing synthetic pathways^14^, and development of theory about the operation of metabolism^15^. Therefore, quantitative thermodynamic data for metabolic reactions at the genome-scale is indispensable in pursuing a systematic and quantitative understanding of metabolism^16^.

Despite the importance of thermodynamics in studying metabolism, its application is still hindered by the lack of accessible experimental data about the thermodynamics of metabolic reactions at the genome-scale. While the most up-to-date human genome-scale metabolic model Recon3D contains about 4,000 metabolites and over 10,000 metabolic reactions^8^, experimentally measured thermodynamic parameters in public databases such as the Thermodynamics of Enzyme-Catalyzed Reactions Database (TECRDB)^17^ and other databases for standard Gibbs energy of formation^18–20^ are only available for a few hundreds of reactions and metabolites, thereby greatly limiting the scope of study on thermodynamics of metabolism.

Several computational methods for prediction of standard Gibbs free energy of metabolites and metabolic reactions have been developed based on the additivity rules for the estimation of molecular properties, i.e., approximation of molecular features by summing up different parts of a molecule^21^. Among those methods, the most noteworthy one is the group contribution (GC) method, which relies on an expert-defined set of chemical groups to decompose the structure of molecules and estimates the standard Gibbs free energy using a linear regression model. The GC method has been applied in predicting standard Gibbs free energy of molecules and reactions in gas phase^21–23^, liquid and solid phase^24^, and biochemical reactions in aqueous solutions^25^. Several algorithms such as the component contribution (CC) method^26^, fingerprint contribution (FC) method^27^, and automated fragmentation method^28^ have been developed to improve the accuracy of prediction^26, 29–31^ and bypass the requirement of expert-defined chemical groups^27, 28^. Nevertheless, the scope of prediction is still limited to metabolites that only contain chemical groups included in molecules in the training set. For example, eQuilibrator 3.0, the most up-to-date database for Gibbs free energy of metabolic reactions predicted using the component contribution method, provides standard Gibbs free energy of about 5,000 human metabolic reactions, covering only one third of the 13,543 reactions in the human genome-scale metabolic model, Recon3D^32^. *Ab initio* quantum chemical computation can calculate standard Gibbs free energy change of biochemical reactions^33–35^ without the need of training datasets, yet the applicability of these methods to metabolites with complex molecular structure is still limited by the extremely high computational cost.

Because of the limitation of existing methods, developing a novel algorithm that can efficiently and accurately predict standard Gibbs free energy of metabolic reactions is an urgent need of the field. The most important reason behind the limitation of the existing methods, almost all of which are derivatives of the GC method, is the dependence of these methods on the ability to decompose the molecular structures in the training set and the molecules to be predicted using the same set of chemical groups. Deep learning models based on graph neural network (GNN) offers a promising possibility to address this problem. Instead of considering a molecule as a linear combination of pre-defined groups, GNN-based models directly treat the molecular structure as a graph and preserve important chemical information at the level of atoms. Hence, GNN-based methods have been continually creating new records on various tasks of predicting properties of molecules^36–40^ with broad applications in chemical property prediction^41^, molecular modelling^42^ and drug discovery^43, 44^. However, to the best of our knowledge, its application for Gibbs free energy prediction has never been reported.

To address these limitations of existing methods and understand the design principles governing thermodynamics of metabolism on genome-scale, we develop a GNN-based machine learning model, dGbyG, for prediction of standard Gibbs free energy change of metabolic reactions from molecular structure of metabolites. We apply two strategies, error randomization and weighing of training data, to improve the accuracy and robustness of our model and offer a quantitative estimation of the uncertainty in the predicted values. An unbiased comparison benchmarking the performance dGbyG and previous models shows that dGbyG greatly outperforms all existing methods in both accuracy and versatility with fewer training data.

In metabolic modeling, flux balance analysis (FBA) with genome-scale models is a standard approach. Under the assumptions of flux balance constraints and the optimization of a metabolic goal such as maximization of biomass production, FBA computes the flux configuration by solving a linear programming problem. A major challenge to FBA is how to effectively constrain the flux space by introducing additional constraints such as those related to thermodynamics.

Interdependence between standard Gibbs energy change, concentrations of metabolites, and direction of a reaction provides a natural point to integrate thermodynamics into genome-scale metabolic models. Several approaches, such as energy balance analysis (EBA)^45^, network- embedded thermodynamic analysis (NET analysis)^46^ and thermodynamics-based metabolic flux analysis (TMFA)^47^ or thermodynamics-based flux balance analysis (TFBA), have been developed to achieve this goal^48^. Previous studies^13, 49, 50^ have shown that thermodynamics-based metabolic modeling has unparalleled advantages in metabolic network reconstruction^9, 10^, prediction of phenotypes^11^ and discovery of biological principles^7^, and so on. Nevertheless, the application of these approaches is greatly limited by the lack of accurate thermodynamics data at the genome scale^32^.

With standard Gibbs free energy values predicted by dGbyG for the human genome-scale metabolic model Recon3D, we find that thermodynamic features of reactions are coupled with topological properties of the network and expressional heterogeneity of the associated metabolic enzymes, indicating that the distribution of thermodynamic parameters within the human metabolic network might be shaped by evolutionary constraints. We also develop an extended TFBA framework to curate the state-of-the-art human GEM, Recon3D, with the assistance of genome-scale thermodynamic data^8^. To our surprise, despite being the most up-to-date genome-scale model for human metabolism, Recon3D still have inaccuracies in reaction directionality and chemical equations, highlighting the potential for thermodynamic data to refine and enhance metabolic modeling. These findings not only elucidate the design principles of metabolic networks that connect thermodynamics, network topology, and enzyme abundance, but also offer practical frameworks for the thermodynamics-based curation of genome-scale metabolic networks.

## Results

### Development of dGbyG

Experimental data of standard Gibbs free energy for metabolites and metabolic reactions used for training the model were obtained from two sources, NIST TECR database^17^ and a table of standard Gibbs free energy of formation of a set of molecules^26^. To generate a high-quality training dataset robust to variability across different measurements, we conducted a series of pre-processing steps (Fig. 1a), in which we removed low-quality data points, curated unbalanced reactions, and standardized the experimental conditions for the measurements (Methods, Supplementary Table 1). To assess the quality of the resulting training set, we computed the standard deviation (SD) of different measurements for the same reaction or the same metabolite and found that the variation across measurements was substantially reduced by the pre- processing procedures (median SD = 1.3kJ/mol in the pre-processed data compared to 1.9 kJ/mol in the raw data, Wilcoxon’s rank-sum p = 0.0002, Supplementary Fig. 1). Finally, we computed the average values over different measurements of the same reaction or metabolite, thereby generating 675 values of standard Gibbs free energy for 453 unique reactions and 641 unique metabolites, which were used as the training set.

**Figure 1.**
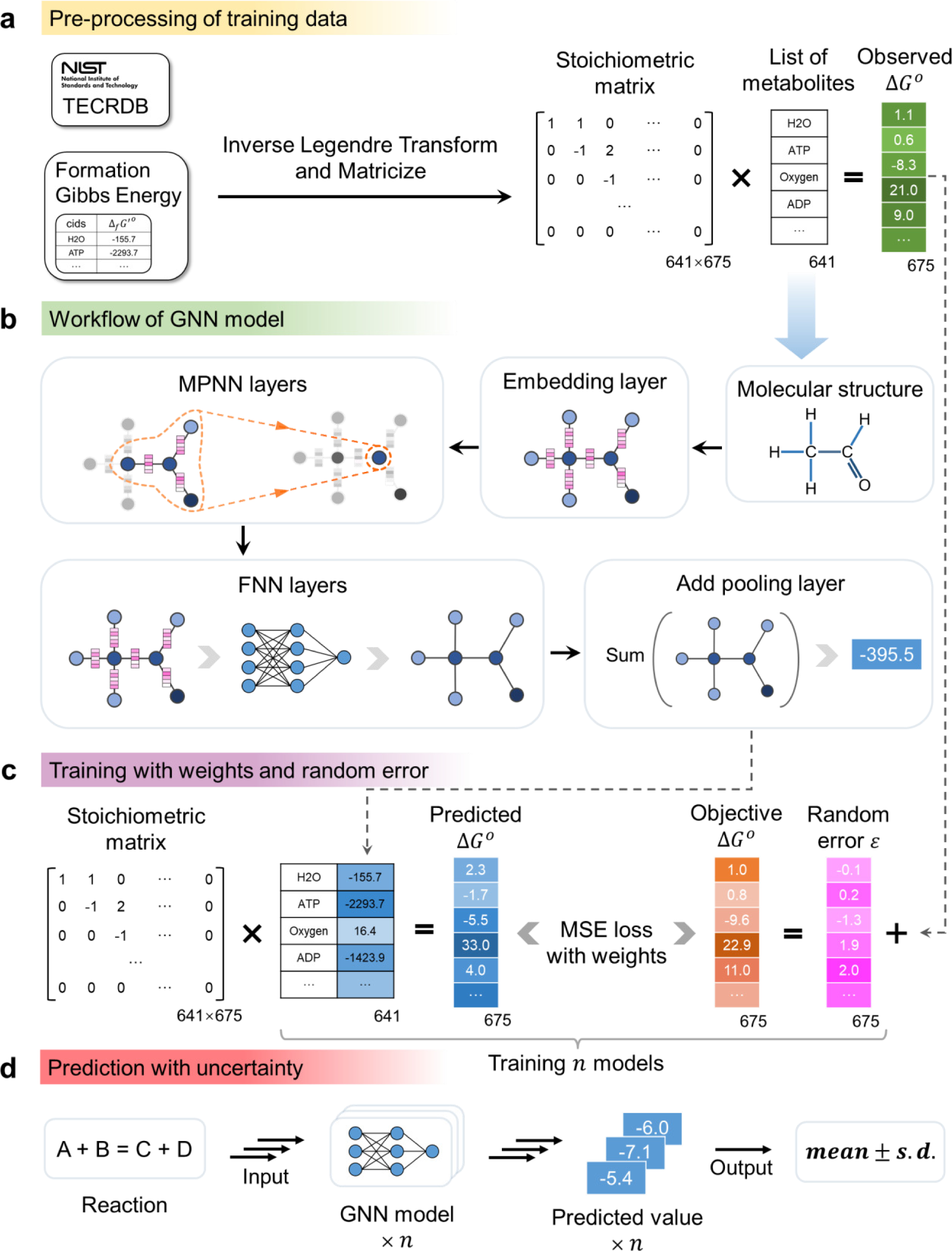
Development of dGbyG. (a) Workflow for pre-processing the training data including 675 experimentally measured standard Gibbs free energy changes for 641 unique metabolites. (b) The architecture of the GNN model used in dGbyG. (c) The method for error randomization and data weighing used in training the model. (d) The method for estimating the uncertainty in predictions.

We developed a GNN-based model dGbyG to predict the standard Gibbs free energy of formation (Δ*_f_G*°) for a metabolite from its molecular structure and estimate the uncertainty in the predictions. We converted the topological structure of molecules to a molecular graph, in which nodes represent atoms and edges represent bonds in the molecule, and parameterized the nodes and edges using chemical features of the atoms and bonds (Supplementary Table 2). An embedding layer, two message passing (MPNN) layers, three feedforward neural network (FNN) layers, and a pooling layer (Fig. 1b) were then included in the GNN model, accepting the molecular graph as the input and computing the Δ*_f_G*° of the molecule by summing up features of all nodes in the pooling layer. Standard Gibbs free energy change of metabolic reactions, Δ*_r_G*°, were then computed from the weighted sum of the Δ*_f_G*° of the substrates and products with the stoichiometric coefficients as the weights (Methods).

To estimate the uncertainty in the model prediction caused by noises in the training set and parameter uncertainty of the model, we developed a bootstrapping-based error randomization approach. Briefly, we assumed a Gaussian distribution for the experimental measurements of the standard Gibbs free energy of each reaction and metabolite, estimated the mean and standard deviation of those distributions from the training data, and then randomly sampled 100 data points from those distributions and used them to train the model separately (Fig. 1c). We also computed a variability index *α* for each data point in the training set and assigned weights to different terms in the loss function accordingly (Supplementary Fig. 2,3). This procedure resulted in 100 sets of model parameters and relevant predictions, which were finally used to estimate the mean and variance of the predictions (Fig 1d, Methods).

Taken together, our model integrates several approaches to provide a high-quality dataset for model training, allow simultaneous prediction of Δ*_f_G*° and Δ*_r_G*°, and characterize the uncertainty in the model predictions. To facilitate the application of dGbyG to genome-scale metabolic models in various scenarios, we implement dGbyG by a Python package that offers user-friendly modules for prediction with genome-scale metabolic models and accepts a variety of molecular identifications and input formats, including SMILES, InChi, KEGG entry, MetaNetX ID, MOL file, RXN file, and so on.

### dGbyG outperforms previous models in accuracy and versatility

To evaluate the performance of dGbyG, we performed 10-fold cross-validation to assess its accuracy and compared its performance with those of other methods^26–28^. We first tested several metrics for the accuracy of prediction, including Pearson’s correlation coefficient (Pearson’s R), mean absolute error (MAE), and root mean squared error (RMSE). The comparison shows that dGbyG greatly improved the accuracy compared to the previous models including eQuilibrator, dGPredictor, and FC (MAE = 9.18 kJ/mol compared to 16.02 kJ/mol for the best of previous models, RMSE = 25.88 kJ/mol compared to 47.85 kJ/mol for the best of previous models, Pearson’s R = 0.997 compared to 0.994 for the best of previous models, Table 1). We then confirmed that the accuracy of dGbyG has been significantly improved by the introduction of the variability index *α* in the loss function, while the error randomization approach, which gives an estimate of the uncertainty in the prediction, did not impair the accuracy (Fig. 2a). We then compared the distribution of prediction error between dGbyG and other models and again confirmed that dGbyG greatly outperformed all previous models in accuracy (median of absolute prediction error = 4.11 kJ/mol for dGbyG compared to 5.33 kJ/mol for the best of previous models, Mood’s median p-value = 1.16×10^-33^, Figure 2b,c).

**Table 1.** Cross-validation results of dGbyG and other methods

|  | Pearson's R | Mean of absolute error (MAE)<sup>#</sup> | Root mean square error (RMSE)<sup>#</sup> |
| --- | --- | --- | --- |
| eQuilibrator | 0.983 | 19.46 | 82.88 |
| dGPredictor | 0.994 | 16.26 | 47.85 |
| FC | 0.994 | 16.02 | 49.46 |
| **dGbyG** | **0.997** | **9.18** | **25.88** |

The state-of-the-art (SOTA) results are shown in bold.
<sup>#</sup> The unit is kJ/mol

**Figure 2.**
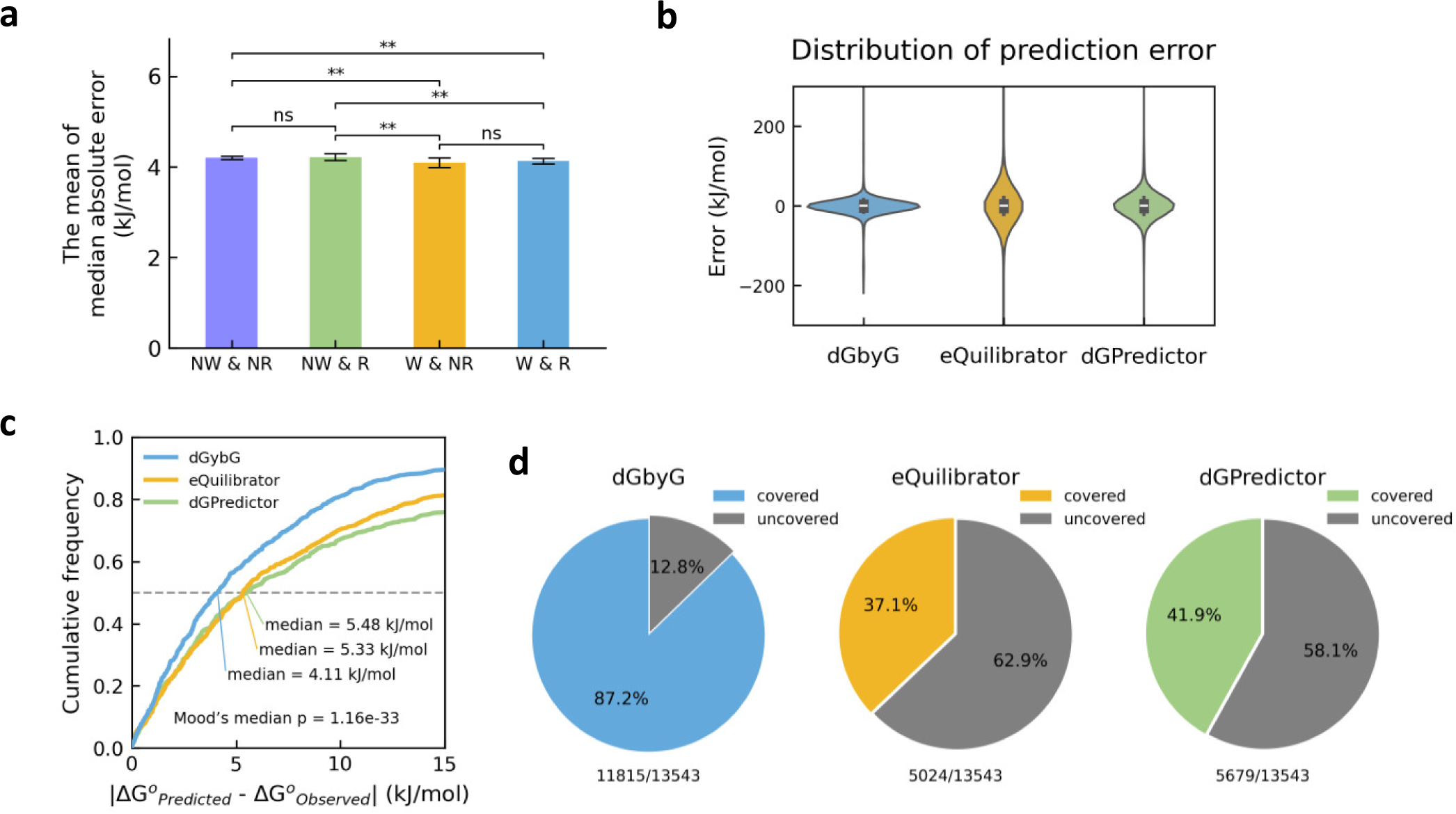
dGbyG outperforms previous methods in accuracy and versatility. (a) Comparison of the mean of median absolute error (MAE) across different combinations of data weighing and error randomization. W: With weighing; R: with error randomization; NW: No weighing; NR: No error randomization. Error bars indicate 95% confidence interval (CI) (n = 10 independent replicates for each condition). ns: no significance (p>=0.05), *: 0.05>p>=0.01, **: p<0.01. (b) Distributions of prediction error of dGbyG, eQuilibrator, and dGPredictor. The prediction error of dGbyG was computed under 10-fold cross-validation, while eQuilibrator and dGPredictor were under leave-one-out cross-validation. (c) Distributions of absolute prediction error of dGbyG, eQuilibrator, and dGPredictor. The cross- validation methods were the same as that in (b). (d) Percentage of reactions in the human genome-scale metabolic network Recon3D for which the standard Gibbs free energy change can be predicted by dGbyG, eQuilibrator, and dGPredictor.

A major limitation to the previous GC-based methods is that they cannot predict the Gibbs free energy for metabolites with chemical groups unseen in the training set. Therefore, we assessed the versatility of dGbyG and other models by testing their ability to make predictions for all reactions included in the human genome-scale metabolic network model Recon3D. We found that dGbyG can predict the Gibbs free energy for almost all reactions in Recon3D (11815 out of 13543 reactions, which represents for 87.2% of all reactions in Recon3D, Figure 2d), which is more than twice of the number of reactions that can be predicted by eQuilibrator (5024 reactions, 37.1%) and dGPredictor (5679 reactions, 41.9%). The reason that prediction by dGbyG is unavailable for a minimal set of reactions is that one or more metabolites involved in those reactions do not have clearly defined molecular structure, hence structure-based predictions of Gibbs free energy for these reactions are impossible. These results together highlight the unprecedented versatility of dGbyG, which is not limited to molecular structures and chemical groups seen in the training data. To the best of our knowledge, dGbyG has achieved the highest accuracy among all existing machine learning models and the highest possible coverage of reactions in genome-scale metabolic models.

### dGbyG outperforms previous models in robustness and generalization ability

We next benchmarked the robustness of dGbyG and other methods by testing the dependence of their accuracy on the size of the training set. We conducted 10-fold cross-validation, 5-fold cross-validation, and 2-fold cross-validation for dGbyG and two previous models eQuilibrator and dGPredictor, which means that the proportion of total available data used for training the model was narrowed down to 90%, 80%, and 50 %, respectively. We found that when the size of training set shrunk, the accuracy of dGbyG was not impaired, while both eQuilibrator and dGPredictor continually lost their predictive power (Pearson’s R in the 2-fold cross validation = 0.997 for dGbyG compared to -0.075 for eQuilibrator and -0.082 for dGPredictor, Figure 3a). In fact, using only half of the available data for training, dGbyG still outperformed the accuracy that eQuilibrator and dGPredictor achieved using all available data for training the model, highlighting its robustness in the situation of few training data.

**Figure 3.**
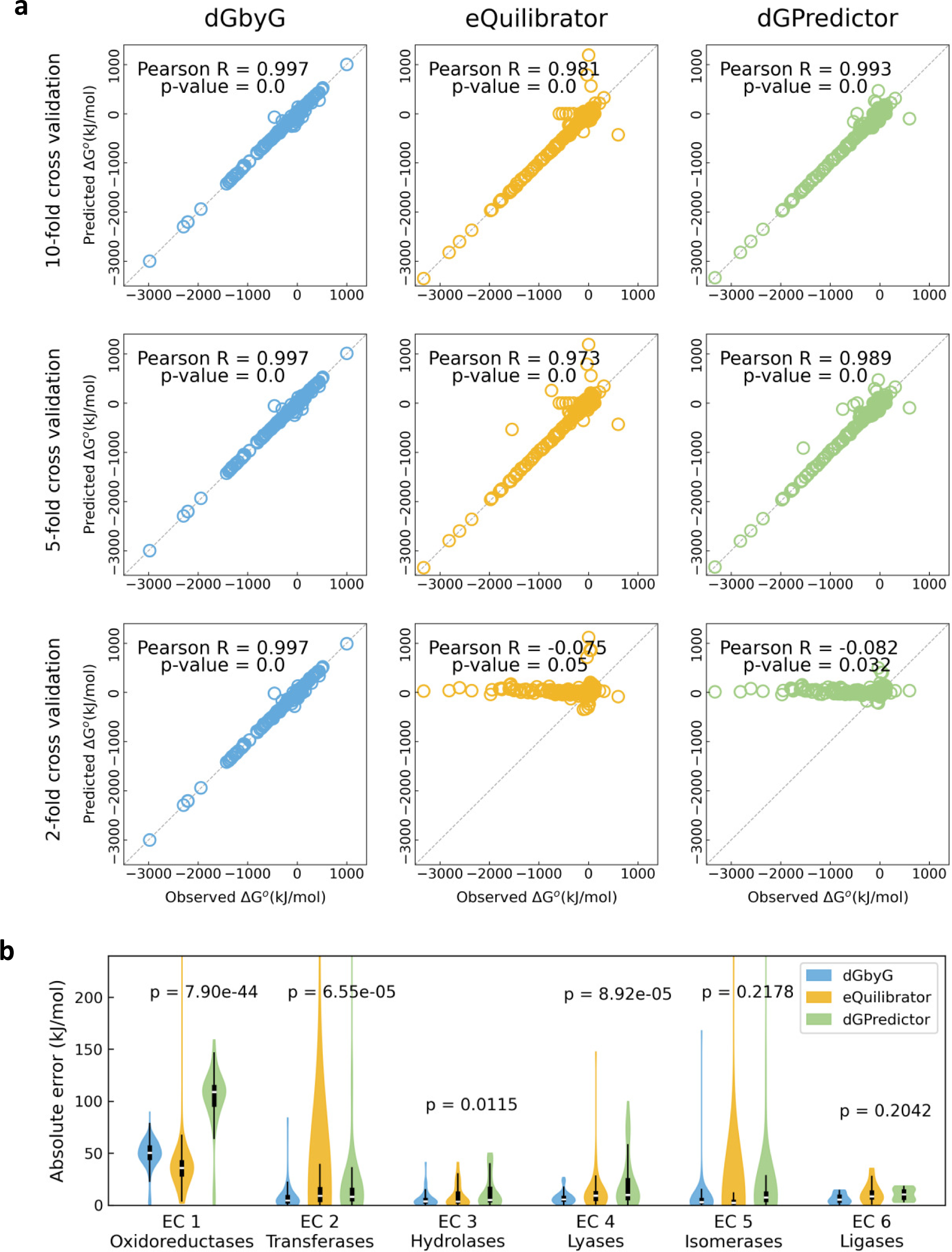
dGbyG outperforms previous methods in robustness and generalization ability. (a) Scatter plots comparing actual and model-predicted standard Gibbs free energy change for dGbyG (left), eQuilibrator (middle), and dGPredictor (right) under 10-fold (top), 5-fold (middle), and 2-fold (bottom) cross-validation. (b) Distributions of prediction error of dGbyG, eQuilibrator, and dGPredictor under leave-one-group-out cross-validation as classified by enzyme commission (EC) codes. P-values were computed using the Alexander-Govern test.

To test whether dGbyG can generate reliable predictions for reactions with mechanisms unseen in the training set, we further performed leave-one-group-out cross-validation for dGbyG and previous methods, in which the reactions were categorized into six groups according to their mechanisms defined by the enzyme commission (EC) codes of the associated enzymes (1 for oxidoreductases, 2 for transferases, 3 for hydrolases, 4 for lyases, 5 for isomerases, and 6 for ligases). In each round of the cross-validation, one group of reactions was used as the validation set, while reactions in the other groups were used to train the model. Amazingly, the accuracy of dGbyG in such cross-validation not only substantially exceeded those of eQuilibrator and dGPredictor, but also became comparable to that of *ab initio* quantum chemistry computations based on the density functional theory^33^ (DFT), which directly models the electronic structure of the atoms and bonds using principles of quantum mechanics without the need of training set and is believed to be the most reliable approach for molecular modeling (dGbyG resulted in lower MAE than that of DFT in three out of six groups, Table 2). However, DFT-based predictions are only available for a minimal set consisting of 150 reactions, while for eQuilibrator and dGPredictor, the removal of reactions with the same mechanism from the training set greatly impaired the accuracy. These results demonstrate that dGbyG outperforms previous GC-based machine learning methods in generalization ability, which allows dGbyG to achieve accuracy comparable to that of first-principle quantum chemistry without requiring training data with similar reaction mechanisms.

**Table 2.** MAE in leave-one-group-out cross-validation for dGbyG and other methods

|  | EC 1 | EC 2 | EC 3 | EC 4 | EC 5 | EC 6 |
| --- | --- | --- | --- | --- | --- | --- |
| eQuilibrator | 38.09 | 60.20 | 8.95 | 13.01 | 21.12 | 12.05 |
| dGPredictor | 102.29 | 26.14 | 13.50 | 21.91 | 19.76 | 10.41 |
| DFT | **7.95** | **8.03** | 6.49 | 7.82 | **5.48** | 13.31 |
| **dGbyG** | 48.71 | 9.19 | **5.96** | **7.33** | 7.18 | **6.49** |

The SOTA results are shown in bold.
The unit is kJ/mol

### Oxygen is a key determinant of thermodynamic drivers in metabolism

The accuracy, robustness, and versatility of dGbyG make it a promising tool to characterize the thermodynamic properties of metabolic networks at the genome-scale. Hence, we applied dGbyG to predict standard Gibbs free energy change of metabolic reactions in the human genome-scale metabolic network model Recon3D^8^. We first studied the distribution of standard Gibbs free energy of reactions in the model, which clearly showed a bimodal distribution with one large peak near 0 kJ/mol and a minor one located near -400 kJ/mol (Fig. 4a), suggesting that although most metabolic reactions operate close to the thermodynamic equilibrium (i.e. Δ*_r_G*° ≈ 0 kJ/mol), a small subset of reactions are far from equilibrium (i.e. Δ*_r_G*° ≪ 0 kJ/mol) and might serve as the thermodynamic driving force of metabolic processes. We refer to that set of reactions with Δ*_r_G*° < −300 kJ/mol as thermodynamic driver reactions (TDRs) in the following discussions.

To characterize the features of thermodynamic driver reactions, we first investigated the substrates and products participating in these reactions. It has been reported in previous studies with a small set of NAD(P)-related oxidation-reduction reactions that the oxygen-dependent ones tend to have larger negative Δ*_r_G*° compared to the oxygen-independent ones, but whether this rule applies to reactions with all possible mechanisms remains unknown^51^. We first compared the distribution of Δ*_r_G*° in reactions catalyzed by different types of enzymes, and found that the thermodynamic driver reactions, indicated by the minor peak in the distribution near -400 kJ/mol, only appear in reactions catalyzed by oxidoreductases (Fig. 4b). Furthermore, to identify key metabolites participating in these thermodynamic driver reactions, we selected metabolites that appear in more than 100 reactions and ranked them by the median absolute Δ*_r_G*° of the reactions associated with that metabolite (Fig. 4c), showing that the oxygen, NADPH, NADP, and hydrogen peroxide are key participants of these thermodynamic driver reactions. We next counted the numbers of thermodynamic driver reactions involving oxygen, NADP, NADPH, and hydrogen peroxide, respectively. We found that although NADP/NADPH are frequently included in thermodynamic driver reactions (400 out of 619 reactions), almost all NADP/NADPH-coupled thermodynamic driver reactions also require oxygen (399 out of 400, Fig. 4d). Interestingly, although hydrogen peroxide is also involved in a small set of thermodynamic driver reactions (17 reactions, Fig. 4d), reactions involving both oxygen and hydrogen peroxide are closer to thermodynamic equilibrium compared to those reactions involving oxygen but not hydrogen peroxide (Fig. 4e). These results together indicate that oxygen is the key determinant of thermodynamic driver reactions, rendering these reactions thermodynamically irreversible and potentially rate-limiting steps controlling metabolic fluxes through a pathway^5^.

**Figure 4.**
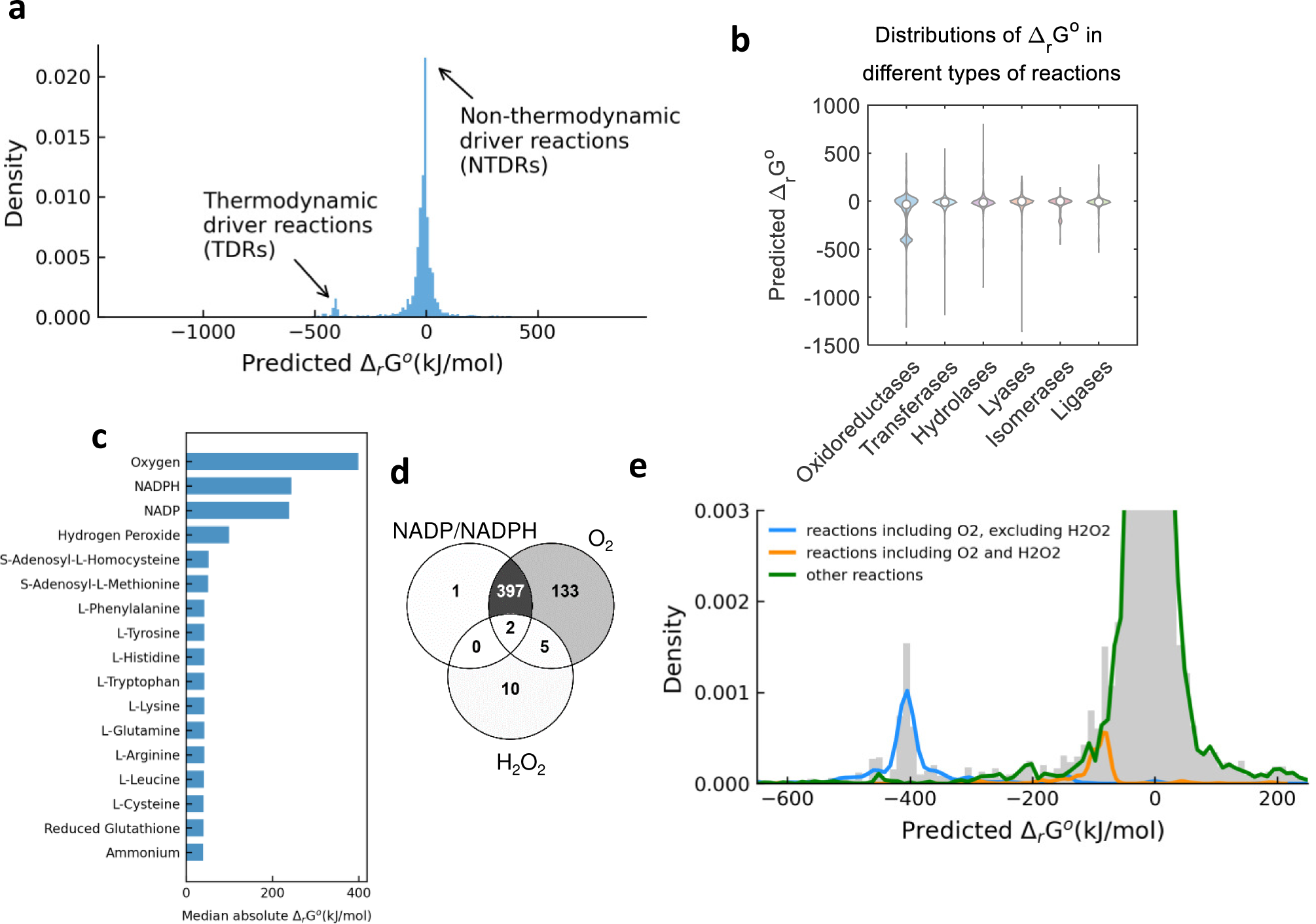
Thermodynamic features of reactions in human metabolism. (a) Distribution of standard Gibbs free energy change for reactions in the human genome-scale metabolic network Recon3D. (b) Violin plots comparing the distributions of standard Gibbs free energy change for reactions catalyzed by different categories of enzymes. (c) Median values of absolute standard Gibbs free energy change in reactions involving different metabolites. (d) Number of thermodynamic driver reactions involving oxygen, hydrogen peroxide, and NADP/NADPH. (e) Distributions of standard Gibbs free energy change in reactions involving different combinations of oxygen and hydrogen peroxide.

### Network topological and proteomic features of thermodynamic driver reactions

Our previous work^5^ and other related studies^6^ have theoretically demonstrated that those thermodynamic driver reactions with substantial negative Gibbs free energy change tend to become the rate-limiting step of a linear metabolic pathway, thereby playing important role in regulating local metabolic fluxes. Studies based on control theory have also reported that, to effectively control a dynamic network to evolve to a desired state, the selection of driver nodes, i.e. a minimal set of nodes in the network that is perturbed to control the dynamics of the entire system, largely depends the topology of the network^52, 53^. It is thus tempting to speculate that, for efficient control of metabolic fluxes, thermodynamic driver reactions should have specific topological features distinct from that of non-thermodynamic-driver reactions in the genome-scale metabolic network. To test this hypothesis, we analyzed various network topological features of the thermodynamic driver reactions. Because a genome-scale metabolic network is a bipartite graph including two types of nodes corresponding to reactions and metabolites, we converted it into a unipartite graph which only includes reactions as the nodes, which we named the reaction connectivity map (Fig. 5a, Methods). An edge connecting two reactions in the reaction connectivity map means that velocities of the two reactions are coupled by at least one metabolite that is involved in both reactions. We first compared several metrics for node importance between the thermodynamic driver reactions and other reactions, including node degree (Fig. 5b), closeness (Supplementary Fig. 4a), and betweenness (Supplementary Fig. 4b), and found that thermodynamic driver reactions tend to have significantly lower degree (Wilcoxon’s rank-sum p = 4.41×10^-24^, Fig. 5b) as well as the other two node importance metrics (Wilcoxon’s rank-sum p = 5.71×10^-25^ for closeness and 9.52×10^-7^ for betweenness, Supplementary Fig. 4a,b). It is worth noting that driver nodes identified solely based on network topology using control theory also tend to avoid the high-degree hubs in the network^52^, suggesting that thermodynamic driver reactions identified based on Gibbs free energy share similar topological features of the driver nodes predicted using control theory.

**Figure 5.**
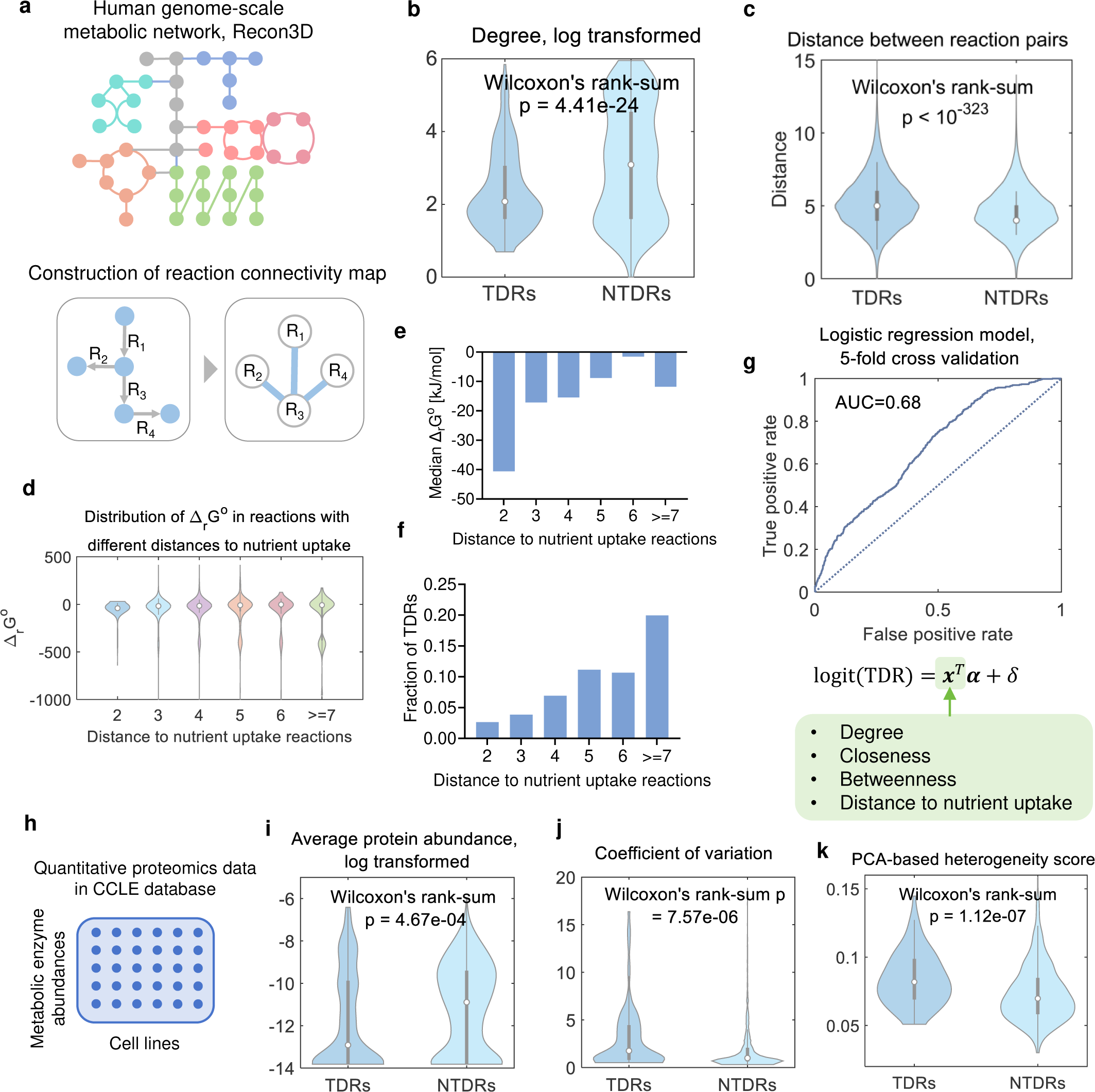
Thermodynamic driver reactions have distinguished network topological features. (a) Construction of the reaction connectivity map from a metabolic network. (b) Violin plots comparing the distributions of log-transformed node degree between thermodynamic driver reactions (TDRs) and other reactions (non-thermodynamic-driver reactions, NTDRs). (c) Violin plots comparing the distributions of distance between two reactions for TDRs and NTDRs. (d) Violin plots comparing distributions of standard Gibbs free energy change in reactions with different distances to the nutrient uptake reactions. (e) Median values of standard Gibbs free energy change in reactions with different distances to the nutrient uptake reactions. (f) Fractions of TDRs in reactions with different distances to the nutrient uptake reactions. (g) Receiver operating characteristics (ROC) curve showing the performance of the logistic regression model predicting TDRs from the network topological features of reactions. (h) Structure of the quantitative proteomics data for human cancer cell lines in the CCLE database. (i) Violin plots comparing log-transformed enzyme abundances between TDRs and NTDRs. (j) Violin plots comparing coefficient of variation in enzyme abundances between TDRs and NTDRs. (k) Violin plots comparing the PCA-based heterogeneity scores of enzyme abundance between TDRs and NTDRs.

We next compared the distributions of Δ*_r_G*° across major metabolic pathways and found that most metabolic pathways contain at least one thermodynamic driver reaction (10 out of 13, Supplementary Fig. 4c), indicating that thermodynamic driver reactions are distributed in different parts of the genome-scale metabolic network and might avoid clustering. To test this hypothesis, we computed the pairwise distance between two thermodynamic driver reactions or between two non-thermodynamic driver reactions in the reaction connectivity map and found that the distance between two thermodynamic driver reactions is significantly longer than that between two non-thermodynamic-driver reactions (Wilcoxon’s rank-sum p < 10^-323^, Fig. 5c), confirming that thermodynamic driver reactions tend to avoid clustering together. We then investigated the locations of thermodynamic driver reactions in the network by correlating Δ*_r_G*° of reactions with their distance to nutrient uptake reactions (Fig. 5d). A reaction with shorter distance to nutrient uptake reactions is more likely located upstream of a metabolic pathway. An interesting finding is that although the upstream reactions with shorter distances to nutrient uptake reactions tend to have larger magnitude negative Δ*_r_G*°(Fig. 5e), the fraction of thermodynamic driver reaction is higher in downstream reactions compared to the upstream ones (Fig. 5f).

These results emphasize that the positioning of thermodynamic driver reactions within the genome-scale metabolic network is not random, potentially enhancing their ability to regulate the metabolic activity of a cell. To further validate this, we built a logistic regression model that can successfully predict whether a metabolic reaction is a thermodynamic driver reaction solely from four variables describing its topological features, including node degree, betweenness, closeness, and distance to nutrient uptake reactions (5-fold cross-validation AUC = 0.68, Fig. 5g). Finally, to confirm that abundances of the enzymes catalyzing these thermodynamic driver reactions can determine the metabolic phenotype of a cell, we obtained quantitative proteomics datasets for 378 human cancer cell lines from the CCLE database^54^ (Fig. 5h). We found that although enzymes catalyzing thermodynamic driver reactions have lower abundance compared to other reactions (Wilcoxon’s rank-sum p = 4.67×10^-4^, Fig. 5i), these enzymes have higher coefficients of variation (CV) across cell lines (Wilcoxon’s rank-sum p = 7.57×10^-6^, Fig. 5j), indicating that their expression is highly dynamic across conditions. We then computed a PCA-based score of heterogeneity to quantify the contribution of each enzyme to the metabolic heterogeneity of cancer cell lines^55^, and found that the variation in expression of enzymes catalyzing thermodynamic driver reactions indeed dominates the metabolic heterogeneity (Wilcoxon’s rank-sum p = 1.12×10^-7^ for the comparison of heterogeneity score between thermodynamic driver reactions and other reactions, Fig. 5k). Taken together, by integrating reaction thermodynamics, network topology and omics data, we have shown that thermodynamic driver reactions with substantial negative values of Δ*_r_G*° have important roles in determining the metabolic phenotypes of cells.

### Curation of genome-scale metabolic models with thermodynamics-based flux balance analysis

The Gibbs free energy change determines whether a process, such as a metabolic reaction, can occur through a specific direction. For a metabolic reaction, its Gibbs free energy change is determined by the standard Gibbs free energy change, Δ*_r_G*°, and concentrations of the substrates and products. Therefore, by integrating standard Gibbs free energy change of metabolic reactions with a genome-scale metabolic network, additional thermodynamic constraints on the flux configurations can be introduced^56, 57^ to address the limitations of the flux balance analysis (FBA) method which often results in non-unique optimal solutions. An important application of this approach is to assign reversibility and directionality of reactions during reconstruction of genome-scale metabolic models^58^. Its accuracy is limited by the scope and quality of available thermodynamic data for metabolic reactions, but such limitation can be addressed by the thermodynamic data predicted by dGbyG at the genome-scale.

Here we developed a thermodynamic-based flux balance analysis (TFBA) approach which introduces two types of constraints: a set of thermodynamic constraints that link the reaction Gibbs free energy change and directionality of fluxes, and an upper limit constraint on the total concentration of metabolites in each compartment of the cell (Methods). We applied TFBA to predict the reversibility and directionality of reactions in Recon3D using dGbyG-predicted values of Δ*_r_G*° and compared the predictions with built-in labels of reaction directionality in Recon3D. We found that for 12 reactions, their directionalities in the original Recon3D model were inconsistent with predictions by TFBA, suggesting that the directions of these reactions originally labeled in Recon3D are thermodynamically infeasible (Fig. 6a, Supplementary Table 3). We then examined these reactions individually and found that for seven of these reactions, their directions in the Recon3D contradicted with literature-based evidence, therefore Recon3D might have mislabeled the directionality of these reactions (Fig. 6b). For example, these reactions include the production of prostaglandin F2alpha (PGF_2α_) from prostaglandin H2 (PGH_2_) in arachidonic acid metabolism, a process known to occur in many different types of human cells such as endothelial cells and adipocytes^59, 60^. While the directionality in Recon3D contradicts with those evidence in literature, TFBA correctly predicted that the reaction should happen in the direction of producing PGF_2α_ from PGH_2_. For some other reactions, their originally assigned directionality in Recon3D results in *de novo* production of oxygen, which is only known to occur through photosynthesis in non-human species such as plants. These reactions can be corrected by reversing their original labels of directionality.

**Fig. 6.**
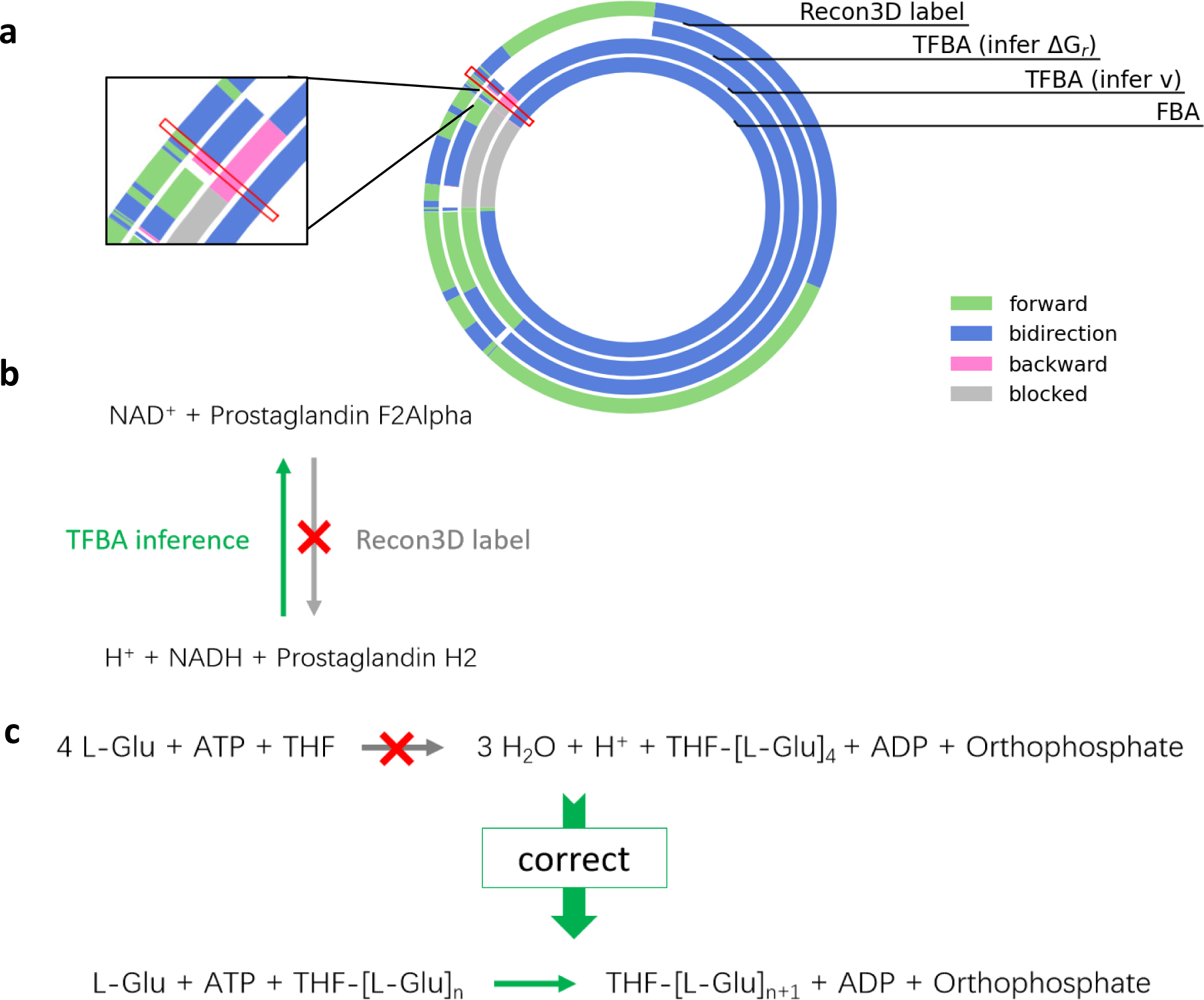
Curation of GEMs with thermodynamics. (a) Comparison between directionalities of reactions in Recon3D predicted by thermodynamics-based flux balanced analysis (TFBA), flux balance analysis (FBA), and their original labels of directionality in the model. (b) An example of a reaction with a potentially mislabeled directionality. (c) An example of a reaction with a potentially mistaken chemical equation.

Some other reactions predicted to be infeasible by TFBA might have improper stoichiometric coefficients, substrates, and products in their chemical equations, which can be addressed by correcting the chemical equation. For example, predictions by TFBA suggested that a reaction in Recon3D which adds four glutamate molecules onto 5,6,7,8-tetrahydrofolate (THF) to form THF- glutamate_4_ (THF-[Glu]_4_) while consuming only one ATP is thermodynamically infeasible.

Experimental evidence in literatures suggest that the formation of poly-glutamate derivatives of THF consists of multiple reaction steps, in which a single ATP molecule is consumed in each step^61, 62^. Thereby, correction of the chemical equation of that reaction can address the thermodynamic infeasibility (Fig. 6c). Although TFBA cannot directly predict the correct chemical equations for all inaccuracies in a genome-scale metabolic model, it can be used to identify potential thermodynamically infeasible reactions, therefore guiding the manual curation of those models.

## Discussion

In this study, we develop a graph neural network model, dGbyG, for predicting thermodynamic features of metabolic reactions at the genome-scale. The model has several remarkable advantages compared to previous models: first, the application of graph neural networks in modeling the molecular structures of metabolites enables the model to efficiently extract information at the levels of atoms and bonds and make accurate predictions for metabolites with chemical groups unseen in the training data and metabolic reactions of mechanisms distinct from those in the training data. Second, with the strategies of error randomization and data weighing that address the uncertainty in the training data and predicted values, we were able to further improve the robustness of our model and give an estimation of the uncertainty in the prediction without impairing the accuracy. Finally, to enhance the usability of our algorithm to other researchers in the field, we implemented our GNN model in a python package dGbyG that encapsulates all necessary steps for the prediction of Gibbs free energy from the chemical equation of a reaction and accepts a variety of different molecular IDs and formats as the input. The comparison with previous methods demonstrates that dGbyG has achieved state-of-the-art performance in accuracy, robustness, versatility, and generalization ability.

Given the versatility of dGbyG, we were able to predict thermodynamic features of metabolic reactions at the genome-scale for the first time. Based on the thermodynamic parameters predicted by dGbyG, many important questions regarding the design principles of metabolic networks can be answered. By studying the distribution of Gibbs free energy over different reactions in a human genome-scale metabolic network, we have shown that there exists a small subset of thermodynamic driver reactions with substantial negative values for the standard Gibbs free energy change. Further investigation of the relationship between reaction thermodynamics, network topology, and heterogeneity in the abundance of metabolic enzymes shows that, these thermodynamic driver reactions, many of which involve oxygen as the substrate, have topological features that are distinct from other reactions, exist in almost all aspects of cellular metabolism, and dominate the metabolic heterogeneity of human cancer cells through the high variability in the abundance of enzymes catalyzing them. These findings suggest that in a metabolic network, there are thermodynamic constraints that govern the location of thermodynamic drivers in the network, potentially for efficient control of metabolic fluxes through these reactions. Further theoretical work is therefore needed to derive the relationship between locations of the thermodynamic driver reactions in a metabolic network and the overall controllability and efficiency of the network.

Another interesting question is whether the relationship between reaction thermodynamics, oxygen, network topology and variability in enzyme abundance is conserved across phylogenetic lineages. Although anaerobic organisms do not use oxygen in their metabolism, they have alternative electron acceptors such as sulfate, nitrate, and sulfur^63^. Because these molecules have lower reduction potential compared to oxygen, reactions using them as electron acceptors will be less strongly driven away from the thermodynamic equilibrium compared to thermodynamic driver reactions we have found in human metabolism, resulting in a different distribution of Δ*_r_G*° over their metabolic networks. How such difference in the global pattern of thermodynamics affects the structure of metabolic networks in anaerobic species also needs further investigation.

Furthermore, by integrating the thermodynamic features of metabolites and reactions predicted by dGbyG with genome-scale metabolic networks using an extended TFBA approach, we were able to identify mislabeled reaction directionalities and chemical equations of reactions in genome-scale reconstruction of metabolic networks. Since the reconstruction of high quality genome-scale metabolic networks is crucial for a quantitative understanding of metabolism, we believe that the application of our approach could greatly benefit the community.

Although dGbyG has superior performance compared to previous methods, it is worth noting that some common challenges to the prediction of Gibbs energy from molecular structures remain. For example, because most of the experimental measurements were made at room temperature, the predictions of dGbyG were also fixed at the room temperature of 298.15 K (25 °C). Therefore, the application of dGbyG to species residing under other temperatures, such as thermophilic bacteria, requires additional procedures to account for the difference in temperature, such as the application of *ab initio* quantum chemical modeling^64^. In addition, three-dimensional molecular conformation also affects the thermodynamic properties of metabolites, but this information is unavailable for most metabolites, thereby limiting the space of increasing the accuracy solely through improving the deep learning model.

Overall, our work offers a powerful computational tool to obtain high-quality thermodynamic data for metabolic reactions at the genome-scale, illustrates the possibility to discover design principles of metabolic networks about the relationship between thermodynamics and network topology, and develops a framework to improve the quality of genome-scale metabolic network reconstructions through the curation with thermodynamic data.

## Methods

### Pre-processing of training data

We first mapped different types of molecular identifiers to SMILES formula, and then removed reactions without clearly defined chemical equations and data points measured under unknown experimental conditions. Based on the pKa values of the molecules predicted by ChemAxon (https://www.chemaxon.com), we then applied the inverse Legendre transformation^65^ to compute the standard Gibbs free energy of the reactions under the same condition of pH=7.0, T = 298.15 K, ionic strength = 0.25 mol/L, and pMg = 14.0. We also developed an algorithm to balance the reactions by automatically adding water molecules and protons to either side of the unbalanced reactions, and then discarded records of standard Gibbs free energy of reactions that cannot be balanced using this procedure. We then computed the mean of standard Gibbs free energy change averaged over different measurements of each reaction, *Δ_r_G*°_Observed_, and the mean of standard formation Gibbs free energy averaged over different measurements of each molecule, *Δ_f_G*°_Observed_. The mean values were used as the training set.

### Structure of GNN model

Molecular structures of metabolites were first converted to SMILES or InChi strings. We also retrieved the corresponding MOL files based on their IDs if needed. We then used RDKit (https://www.rdkit.org/) to process and standardize the molecules by adding missing hydrogen atoms, and converted all structures to molecular graphs with atoms as nodes and bonds as edges, recording atomic number, hybridization, aromaticity, and charge of atoms as node features and type of bonds as edge features. The neural network model starts with the one-hot encoding of the molecular graph, transforming it into the latent space by embedding layers. Then two message passing neural network (MPNN) layers propagate and aggregate the message from bonds and neighboring atoms for each atom together with a residual of its own features, producing the features of each atom in the next layer. Three feedforward neural network (FNN) layers were then used to transform the feature vector of each atom to a single value, followed by an additive pooling layer that sums up all single values of atoms to compute the Δ*_f_G*° as the final output of the neural network. Finally, we compute the standard Gibbs free energy change for a metabolic reaction with the following chemical equation from the model-predicted values of Δ*_f_G*° of the substrates {*S_i_*} and products {*P_i_*} and the stoichiometric coefficients {*a_i_*} and {*b_i_*}:

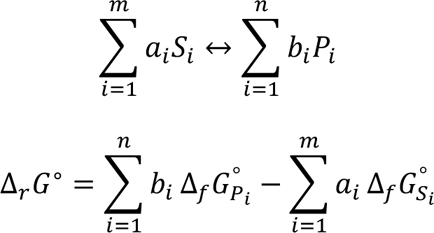

The whole framework is implemented based on PyTorch^66^ and PyG^67^.

### Estimation of uncertainty in the prediction

To define a metric to quantify the variability in experimental measurements of Gibbs free energy, we quantified the relationship between its standard deviation (SD), standard error of the mean (SEM), absolute value, and sample size (i.e. number of different measurements) for each reaction. We did not observe strong correlation between the standard deviation (SD) of experimentally measured Gibbs energy values and other data features (Supplementary Fig. 2a,b), but the significant positive correlation between the standard error of the mean (SEM) and number of measurements suggests that the variability in the mean values decreases as the sample size increases, in other words, measurements with lower SEM are more reliable than the other ones (Supplementary Fig. 2c). Hence, we define an index *α* to assess the variability in the experimental measurements of standard Gibbs energy for each reaction:

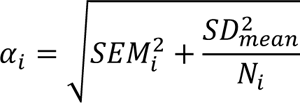

where *SEM_i_* is the SEM of the *i*-th reaction, *SD_mean_* is the mean SD of all reactions in the training set, *N_i_* is the number of different measurements for the *i*-th reaction. The design of *α* aims to stabilize the variability in the data point by approximating it according to the average SD of all reactions when the sample size *N* is small. Otherwise, when the sample size *N* is large, the variability index converges to the actual SEM of the reaction. Under the assumption that the error in the mean value of experimentally measured standard Gibbs energy for each reaction follows a normal distribution with standard deviation *α* equals to the variability index defined above, we randomly sampled 100 error terms from that distribution for each reaction and added it to the original value in the training set to generate 100 different training sets. We then used those 100 training sets to train 100 different networks, resulting in an ensemble of models with different parameters and predictions. The predictions made by 100 networks in the ensemble of models were used to estimate uncertainty in the prediction.

### Topological analysis of metabolic networks

MATLAB .mat file implementing the human GEM Recon3D was retrieved from the online resource http://vmh.life/. An undirected graph named reaction connectivity map was then constructed based on the stoichiometric matrix *S* of the model. In constructing the reaction connectivity map, reactions in Recon3D were treated as nodes, and two reactions were considered to be connected with each other if at least one non-hub metabolite is involved in both reactions. Non-hub metabolites were defined by metabolites that are not ions and molecules that participate in many reactions, such as proton, water, phosphate, NAD/NADH, NADP/NADPH, carbon dioxide, and so on. Degree, closeness and betweenness of each reaction node in the reaction connectivity map were computed using the MATLAB built-in functions degree [] and centrality []. Distances between reactions in the reaction connectivity map were computed using the MATLAB built-in function distances []. Exchange reactions for glucose, glutamine, homoserine, isoleucine, leucine, lysine, methionine, phenylalanine, serine, threonine, tryptophan, tyrosine, and valine were defined as nutrient uptake reactions. The logistic regression model predicting thermodynamic driver reactions from topological features of reactions was built with the degree, betweenness, closeness and distance to nutrient uptake reactions as four independent variables, and the binary label of a reaction being a thermodynamic driver reaction or not as the dependent variable. The model was fit to the dataset containing 6,854 reactions in Recon3D using the MATLAB built-in function glmfit [], and its performance was evaluated by 5-fold cross-validation using the MATLAB function glmval [].

### Analysis of enzyme abundance in human cancer cell lines

Quantitative proteomics data of human cancer cell lines measured by mass spectrometry were retrieved from the non-normalized protein data available at https://gygi.hms.harvard.edu/publications/ccle.html, and proportionally scaled by the sum of intensities of all proteins detected in each cell line. We mapped the gene symbols in the proteomic dataset to Entrez IDs using the online tool SynGO^68^, and computed abundances of enzymes associated with each reaction based on the gene-protein-reaction mapping matrix in Recon3D. We then classified the enzymes into two categories based on their association with the TDRs: one TDR-group consists of enzymes associated with at least one TDR, while the other NTDR-group contains enzymes that are not associated with any TDR. We computed coefficient of variation (CV) for each enzyme, and then performed Z-score normalization of the enzyme abundance matrix so that the distribution of enzyme abundance across cell lines for each reaction has the mean value of zero and the standard deviation of one. Principal component analysis (PCA) was then performed with the Z-score normalized enzyme abundance matrix, and a heterogeneity score for each enzyme was computed by squared sum of the loadings of that enzyme in the top 200 principal components.

### Thermodynamics-based Flux Balance Analysis (TFBA) with upper limit of total metabolite concentration

We develop the framework of TFBA by introducing additional constraints linking fluxes, metabolite concentrations, and reaction thermodynamics to the original mathematical problem of flux balance analysis (FBA). Standard Gibbs energy of reactions predicted by dGbyG were used to parametrize some of these constraints on the flux configurations. Briefly, we incorporated variables corresponding to Δ*_f_G*° of each metabolite, Δ*_r_G*° and Δ*_r_G* of each reaction, concentrations of metabolites, and directionalities of reactions into the original FBA model. A set of nonlinear inequality constraints on Gibbs free energy change of all reactions were then introduced, which require that the net flux carried by the reaction and the reaction Gibbs free energy change Δ*_r_G* = Δ*_r_G*° + *RT* ln *Q* have opposite signs. Δ*_r_G*° was constrained in the range of uncertainty of *Δ_r_G*°_*Predicted*_ for reactions predicted by dGbyG. By introducing a binary variable *a* indicating the direction of the reaction and logarithms of the concentration variables, this nonlinear inequality constraint can be converted to a linear inequality constraint with integer variables, which can be solved under the framework of mixed-integer programming (MIP) problems. To eliminate unrealistic solutions, we also set an upper limit of total concentration of metabolites in each compartment. Because this is a nonlinear constraint on the logarithms of metabolite concentrations, we used a piecewise linear function to approximate it with linear constraints. A detailed description of those constraints is available in Supplementary Information. Subject to these constraints, we determined the upper and lower bounds of the reaction Gibbs free energy change Δ*_r_G* for each reaction by solving the mixed-integer programming (MIP) problems to maximize and minimize Δ*_r_G*. A reaction was classified as reversible if its Δ*_r_G* had a negative lower bound and a positive upper bound. Otherwise, if the upper bound of Δ*_r_G* was negative, the reaction was deemed irreversible and assigned a forward directionality; conversely, if the lower bound was positive, it was assigned a backward directionality. We also computed the upper and lower bounds of the net flux of each reaction to further constrain the directionality of the reactions predicted to be bidirectional based on the range of Δ*_r_G*.

## Data availability

Datasets used for training dGbyG are available at the Gitee repository https://gitee.com/f-wc/dGbyG.git. Standard Gibbs energy change of reactions for Recon3D predicted by dGbyG is available at the Gitee repository https://gitee.com/f-wc/ThermoInfer.git.

## Code availability

Codes and detailed instructions for dGbyG and TFBA are available at the Gitee repositories https://gitee.com/f-wc/dGbyG.git and https://gitee.com/f-wc/ThermoInfer.git.

## Author contributions

W.F. and Z.D. designed the study and wrote the manuscript. W.F. developed the dGbyG and TFBA algorithms and conducted the comparison of performance with previous methods. Z.D. performed the analysis of thermodynamic driver reactions in Recon3D with inputs from D.H. W.Z. analyzed the CCLE proteomics data. C.D. and W.F. curated the mistaken reactions in Recon3D found by TFBA. All authors have read and approved the final manuscript.

## Acknowledgements

The authors thank the National Natural Science Foundation of China (12371489 to ZD) and National Key Research and Development Program of China (2021YFA0911300 and 2021YFF1201000 to ZD) for their generous support. Support for computational resources from the Center for Computational Science and Engineering of Southern University of Science and Technology is gratefully acknowledged.

## References

1. Demetrius, L. Thermodynamics and Evolution. Journal of Theoretical Biology 206, 1–16 (2000).

2. Nelson, D.L., Cox, M.M. & Hoskins, A.A. Lehninger principles of biochemistry, Edn. Eighth edition. (Macmillan Learning, Austin; 2021).

3. Shakhnovich, E. Protein folding thermodynamics and dynamics: where physics, chemistry, and biology meet. Chem Rev 106, 1559–1588 (2006).

4. Weng, C., Faure, A.J., Escobedo, A. & Lehner, B. The energetic and allosteric landscape for KRAS inhibition. Nature (2023).

5. Dai, Z. & Locasale, J.W. Thermodynamic constraints on the regulation of metabolic fluxes. J Biol Chem 293, 19725–19739 (2018).

6. Noor, E. et al. Pathway thermodynamics highlights kinetic obstacles in central metabolism. PLoS Comput Biol 10, e1003483 (2014).

7. Niebel, B., Leupold, S. & Heinemann, M. An upper limit on Gibbs energy dissipation governs cellular metabolism. Nat Metab 1, 125–132 (2019).

8. Brunk, E. et al. Recon3D enables a three-dimensional view of gene variation in human metabolism. Nat Biotechnol 36, 272–281 (2018).

9. Martínez, Verónica S., Quek, L.-E. & Nielsen, Lars K. Network Thermodynamic Curation of Human and Yeast Genome-Scale Metabolic Models. Biophysical Journal 107, 493–503 (2014).

10. Haraldsdóttir, H.S., Thiele, I. & Fleming, R.M.T. Quantitative Assignment of Reaction Directionality in a Multicompartmental Human Metabolic Reconstruction. Biophysical Journal 102, 1703–1711 (2012).

11. Hamilton, Joshua J., Dwivedi, V. & Reed, Jennifer L. Quantitative Assessment of Thermodynamic Constraints on the Solution Space of Genome-Scale Metabolic Models. Biophysical Journal 105, 512–522 (2013).

12. Schultz, A. & Qutub, A.A. Predicting internal cell fluxes at sub-optimal growth. BMC Systems Biology 9 (2015).

13. Kiparissides, A. & Hatzimanikatis, V. Thermodynamics-based Metabolite Sensitivity Analysis in metabolic networks. Metabolic Engineering 39, 117–127 (2017).

14. Sauro, H., Hädicke, O., von Kamp, A., Aydogan, T. & Klamt, S. OptMDFpathway: Identification of metabolic pathways with maximal thermodynamic driving force and its application for analyzing the endogenous CO2 fixation potential of Escherichia coli. PLOS Computational Biology 14 (2018).

15. Mori, M., Cheng, C., Taylor, B.R., Okano, H. & Hwa, T. Functional decomposition of metabolism allows a system-level quantification of fluxes and protein allocation towards specific metabolic functions. Nat Commun 14, 4161 (2023).

16. Ataman, M. & Hatzimanikatis, V. Heading in the right direction: thermodynamics-based network analysis and pathway engineering. Current Opinion in Biotechnology 36, 176–182 (2015).

17. Goldberg, R.N., Tewari, Y.B. & Bhat, T.N. Thermodynamics of enzyme-catalyzed reactions--a database for quantitative biochemistry. Bioinformatics 20, 2874–2877 (2004).

18. Krebs, H.A. & Kornberg, H.L. in Energy Transformations in Living Matter: A Survey. (eds. H.A. Krebs & H.L. Kornberg) 212–298 (Springer Berlin Heidelberg, Berlin, Heidelberg; 1957).

19. Thauer, R.K., Jungermann, K. & Decker, K.H. Energy Conservation in Chemotrophic Anaerobic Bacteria. Bacteriological Reviews 41, 809–809 (1977).

20. Alberty, R.A. Biochemical thermodynamics: applications of Mathematica. Methods Biochem Anal 48, 1–458 (2006).

21. Benson, S.W. & Buss, J.H. Additivity Rules for the Estimation of Molecular Properties. Thermodynamic Properties. The Journal of Chemical Physics 29, 546–572 (1958).

22. Boudart, M. Thermochemical kinetics, 2nd. Ed., Sidney W. Benson, Wiley Interscience, 320 pp., $22.50, New York, 1976. AIChE Journal 23, 613–613 (1977).

23. Ritter, E.R. & Bozzelli, J.W. THERM: THERMODYNAMIC PROPERTY ESTIMATION FOR GAS PHASE RADICALS and MOLECULES. Proceeding of Data For Discovery (1991).

24. Domalski, E.S. & Hearing, E.D. Estimation of the Thermodynamic Properties of Hydrocarbons at 298.15 K. Journal of Physical and Chemical Reference Data 17, 1637–1678 (1988).

25. Mavrovouniotis, M.L., Bayol, P., Lam, T.K.M., Stephanopoulos, G. & Stephanopoulos, G. A group contribution method for the estimation of equilibrium constants for biochemical reactions. Biotechnology Techniques 2, 23–28 (1988).

26. Noor, E., Haraldsdottir, H.S., Milo, R. & Fleming, R.M. Consistent estimation of Gibbs energy using component contributions. PLoS Comput Biol 9, e1003098 (2013).

27. Alazmi, M., Kuwahara, H., Soufan, O., Ding, L. & Gao, X. Systematic selection of chemical fingerprint features improves the Gibbs energy prediction of biochemical reactions. Bioinformatics 35, 2634–2643 (2019).

28. Wang, L., Upadhyay, V. & Maranas, C.D. dGPredictor: Automated fragmentation method for metabolic reaction free energy prediction and de novo pathway design. PLoS Comput Biol 17, e1009448 (2021).

29. Jankowski, M.D., Henry, C.S., Broadbelt, L.J. & Hatzimanikatis, V. Group contribution method for thermodynamic analysis of complex metabolic networks. Biophys J 95, 1487–1499 (2008).

30. Noor, E. et al. An integrated open framework for thermodynamics of reactions that combines accuracy and coverage. Bioinformatics 28, 2037–2044 (2012).

31. Du, B. et al. Temperature-Dependent Estimation of Gibbs Energies Using an Updated Group- Contribution Method. Biophysical Journal 114, 2691–2702 (2018).

32. Beber, M.E. et al. eQuilibrator 3.0: a database solution for thermodynamic constant estimation. Nucleic Acids Res 50, D603–D609 (2022).

33. Joshi, R.P. et al. Quantum Mechanical Methods Predict Accurate Thermodynamics of Biochemical Reactions. ACS Omega 6, 9948–9959 (2021).

34. Jinich, A., et al. Quantum Chemical Approach to Estimating the Thermodynamics of Metabolic Reactions. Scientific Reports 4 (2014).

35. van Speybroeck, V., Gani, R. & Meier, R.J. The calculation of thermodynamic properties of molecules. Chemical Society Reviews 39 (2010).

36. Yang, Q., Ji, H., Lu, H. & Zhang, Z. Prediction of Liquid Chromatographic Retention Time with Graph Neural Networks to Assist in Small Molecule Identification. Anal Chem 93, 2200–2206 (2021).

37. Ma, H. et al. Cross-Dependent Graph Neural Networks for Molecular Property Prediction. Bioinformatics (2022).

38. Fang, X. et al. Geometry-enhanced molecular representation learning for property prediction. Nature Machine Intelligence 4, 127–134 (2022).

39. Wang, Y., Wang, J., Cao, Z. & Barati Farimani, A. Molecular contrastive learning of representations via graph neural networks. Nature Machine Intelligence 4, 279–287 (2022).

40. Fang, Y. et al. Knowledge graph-enhanced molecular contrastive learning with functional prompt. Nature Machine Intelligence 5, 542–553 (2023).

41. Roszak, R., Beker, W., Molga, K. & Grzybowski, B.A. Rapid and Accurate Prediction of pK(a) Values of C-H Acids Using Graph Convolutional Neural Networks. J Am Chem Soc 141, 17142–17149 (2019).

42. Park, C.W. et al. Accurate and scalable graph neural network force field and molecular dynamics with direct force architecture. npj Computational Materials 7, 73 (2021).

43. Xiong, J., Xiong, Z., Chen, K., Jiang, H. & Zheng, M. Graph neural networks for automated de novo drug design. Drug Discovery Today 26, 1382–1393 (2021).

44. Wong, F. et al. Discovery of a structural class of antibiotics with explainable deep learning. Nature (2023).

45. Beard, D.A., Liang, S.-d. & Qian, H. Energy Balance for Analysis of Complex Metabolic Networks. Biophysical Journal 83, 79–86 (2002).

46. Kümmel, A., Panke, S. & Heinemann, M. Putative regulatory sites unraveled by network- embedded thermodynamic analysis of metabolome data. Molecular Systems Biology 2 (2006).

47. Henry, C.S., Broadbelt, L.J. & Hatzimanikatis, V. Thermodynamics-Based Metabolic Flux Analysis. Biophysical Journal 92, 1792–1805 (2007).

48. Soh, K.C. & Hatzimanikatis, V. in Metabolic Flux Analysis: Methods and Protocols. (eds. J.O. Krömer, L.K. Nielsen & L.M. Blank) 49–63 (Springer New York, New York, NY; 2014).

49. Dash, S. et al. Thermodynamic analysis of the pathway for ethanol production from cellobiose in Clostridium thermocellum. Metabolic Engineering 55, 161–169 (2019).

50. Krumholz, E.W. & Libourel, I.G.L. Thermodynamic Constraints Improve Metabolic Networks. Biophysical Journal 113, 679–689 (2017).

51. Goldford, J.E., George, A.B., Flamholz, A.I. & Segre, D. Protein cost minimization promotes the emergence of coenzyme redundancy. Proc Natl Acad Sci U S A 119, e2110787119 (2022).

52. Liu, Y.Y., Slotine, J.J. & Barabasi, A.L. Controllability of complex networks. Nature 473, 167–173 (2011).

53. Basler, G., Nikoloski, Z., Larhlimi, A., Barabasi, A.L. & Liu, Y.Y. Control of fluxes in metabolic networks. Genome Res 26, 956–968 (2016).

54. Nusinow, D.P. et al. Quantitative Proteomics of the Cancer Cell Line Encyclopedia. Cell 180, 387–402 e316 (2020).

55. Xiao, Z., Dai, Z. & Locasale, J.W. Metabolic landscape of the tumor microenvironment at single cell resolution. Nat Commun 10, 3763 (2019).

56. Henry, C.S., Broadbelt, L.J. & Hatzimanikatis, V. Thermodynamics-based metabolic flux analysis. Biophys J 92, 1792–1805 (2007).

57. Kummel, A., Panke, S. & Heinemann, M. Systematic assignment of thermodynamic constraints in metabolic network models. BMC Bioinformatics 7, 512 (2006).

58. Thiele, I. & Palsson, B.O. A protocol for generating a high-quality genome-scale metabolic reconstruction. Nat Protoc 5, 93–121 (2010).

59. Bresson, E., Lacroix-Pepin, N., Boucher-Kovalik, S., Chapdelaine, P. & Fortier, M.A. The Prostaglandin F Synthase Activity of the Human Aldose Reductase AKR1B1 Brings New Lenses to Look at Pathologic Conditions. Front Pharmacol 3, 98 (2012).

60. Michaud, A. et al. Prostaglandin (PG) F2 alpha synthesis in human subcutaneous and omental adipose tissue: modulation by inflammatory cytokines and role of the human aldose reductase AKR1B1. PLoS One 9, e90861 (2014).

61. Tomsho, J.W., Moran, R.G. & Coward, J.K. Concentration-dependent processivity of multiple glutamate ligations catalyzed by folylpoly-gamma-glutamate synthetase. Biochemistry 47, 9040–9050 (2008).

62. Sheng, Y. et al. Mutagenesis of Folylpolyglutamate Synthetase Indicates That Dihydropteroate and Tetrahydrofolate Bind to the Same Site. Biochemistry 47, 2388–2396 (2008).

63. Leng, H., Wang, Y., Zhao, W., Sievert, S.M. & Xiao, X. Identification of a deep-branching thermophilic clade sheds light on early bacterial evolution. Nat Commun 14, 4354 (2023).

64. Hadadi, N., Ataman, M., Hatzimanikatis, V. & Panayiotou, C. Molecular thermodynamics of metabolism: quantum thermochemical calculations for key metabolites. Physical Chemistry Chemical Physics 17, 10438–10453 (2015).

65. Alberty, R.A. Inverse Legendre Transform in Biochemical Thermodynamics: Illustrated with the Last Five Reactions of Glycolysis. The Journal of Physical Chemistry B 106, 6594–6599 (2002).

66. Paszke, A. et al. PyTorch: An Imperative Style, High-Performance Deep Learning Library. (Curran Associates, Inc., 2019).

67. Fey, M. & Lenssen, J.E. Fast Graph Representation Learning with PyTorch Geometric. (2019).

68. Koopmans, F., et al. SynGO: An Evidence-Based, Expert-Curated Knowledge Base for the Synapse. Neuron 103, 217–234 e214 (2019).

