## Supplementary information for "Unraveling principles of thermodynamics for genome-scale metabolic networks using graph neural networks"

### Supplementary Methods

#### TFBA with concentration upper limit

The original method for flux balance analysis (FBA) relies on the hypothesis that for each metabolite in the model, fluxes producing it are balanced by fluxes consuming it. Therefore, we have the following linear constraints on the fluxes in a metabolic network:

$$S \cdot v = 0, \tag{1}$$

$$v_l \le v \le v_u, \tag{2}$$

where $S$ is a $m \times r$ stoichiometric matrix, $v$ is an $r \times 1$ flux vector (dimensionless variables), $m$ is the total number of metabolites and $r$ is the total number of reactions. $v_l$ and $v_u$ are two $r \times 1$ vectors representing the lower and upper bounds of flux for each reaction.

The second law of thermodynamics assert that a reaction can only occur through the direction by which the Gibbs free energy is released, in other words, the Gibbs free energy change of a thermodynamically feasible reaction should be negative. Therefore, non-linear constraints on Gibbs free energy change of each reaction can be introduced as following,

$$a \in \{0,1\}, \tag{3}$$

$$(a-1) \times 1000 \le v \le a \times 1000, \tag{4}$$

$$-a \times M \le \Delta_r G \le (1-a) \times M, \tag{5}$$

$$M = 4000, \tag{6}$$

where $a$ is an $r \times 1$ vector of binary instrumental variables indicating the directionality of reactions, $\Delta_r G$ is an $r \times 1$ vector of Gibbs free energy change of reactions, M is a sufficiently large positive number. Here we set $M$ to 4000 because its magnitude is much larger than the standard Gibbs free energy change of typical metabolic reactions.

In fact, metabolic reactions in cells are almost impossible to reach equilibrium ($\Delta G_r = 0$) stably because a slight change in concentrations of metabolites would break the equilibrium and changes are happened constantly. Plus, when a reaction is very close to equilibrium, its net flux would be few even the enzyme was numerous. Consequently, we set a lower limit of absolute $\Delta_r G$ to eliminate a large $v$ existing in a reaction of near equilibrium. So we replace inequalities (5) by

$$-a \times M + \tau \le \Delta_r G \le (1-a) \times M - \tau, \tag{7}$$

$$\tau = 0.05, \tag{8}$$

where $\tau$ is lower limit of the absolute $\Delta_r G$. We set $\tau = 0.05$ here because it is small enough compared with the uncertainty of predicted standard Gibbs energy change, so that do not affect TFBA on most tasks.

For each reaction, $\Delta_r G$ is a function of its standard Gibbs energy change and concentrations of reactants:

$$\Delta_r G = \Delta_r G^{\circ} + RTln(10)S^T \cdot z, \quad (9)$$

$$z = log_{10}(c), \quad (10)$$

$$z_l \le z \le z_u, \quad (11)$$

where $\Delta_r G^{\circ}$ is a $r \times 1$ vector of standard Gibbs energy change of reactions, $R = 8.314 \times 10^{-3}$ $\mathrm{kJ \cdot mol^{-1} \cdot K^{-1}}$ is the gas constant, $T$ is the temperature, $S^T$ is the transposition of the stoichiometric matrix $S$, $c$ is a $m \times 1$ vector of the concentrations of metabolites, $z$ is the base-10 logarithm of $c$. $z_l$ and $z_u$ are two $m \times 1$ vectors representing for the lower and upper bounds of each element in $z$.

The standard Gibbs free energy change $\Delta_r G^{\circ}$ of most reactions could be predicted by dGbyG or retrieved from experimental data. For those reactions, we computed the confidence intervals of the $\Delta_r G^{\circ}$ values based on the estimation of uncertainty made by dGbyG or experimental data available from multiple sources. According to the confidence intervals, the following constraints were added:

$$\mu_i \le \Delta_r G_i^{\circ} - \Delta_r G_{p,i}^{\circ} \le \upsilon_i, \{i = 1,2, \ldots, n | \Delta_r G_{p,i}^{\circ} \ is \ known\}, \quad (12)$$

where $\Delta_r G_{p,i}^{\circ}$ and $\Delta_r G_i^{\circ}$ are the known values of standard Gibbs energy change available from prediction and experimental measurement, respectively. $\mu_i$ and $\upsilon_i$ are the lower and upper bounds of the error in prediction made by dGbyG or experimental measurements.

It is also worth noting that the $\Delta_r G^{\circ}$ of reactions depends on the standard Gibbs free energy of formation of metabolites involved in these reactions. For this reason, we have linear equality constraints linking these two:

$$\Delta_r G^{\circ} = S^T \cdot \Delta_f G^{\circ}, \quad (13)$$

where $\Delta_f G^{\circ}$ is a $m \times 1$ vector representing for the standard formation Gibbs energy of metabolites. Note that this equation makes sure that each metabolite has a single value of $\Delta_f G^{\circ}$ in the system, thereby allowing the algorithm to accommodate thermodynamic parameters from different sources, even those that may be thermodynamically inconsistent and violate the first law of thermodynamics. This ensures that the entire system is thermodynamically consistent by coupling the values of $\Delta_f G^{\circ}$ and $\Delta_r G^{\circ}$.

Finally, a constraint that sets the upper limit of total concentration of metabolites in each compartment was added to the model:

$$M_{cm} \cdot c \le C_{upper}, \quad (14)$$

where $M_{cm}$ is a $k \times m$ matrix in which $k$ means the number of separated intracellular compartments such as cytosol and mitochondria. $M_{cm\ i,j}$ is 1 if the metabolite $j$ is in compartment $i$, otherwise its value is zero. $C_{upper}$ is a $k \times 1$ vector consisting of the upper bounds in all compartments. This constraint can also be written as a constraint on the logarithms of metabolite concentrations $z$:

$$M_{cm} \cdot 10^{z} \le C_{upper}, \quad (15)$$

Because this constraint is non-linear, the computational cost of solving the resulting optimization problem is extremely high with a genome-scale metabolic network. Therefore, we convert it into linear constraints by approximating the exponential function between $z$ and $c$ using a piecewise linear function:

$$b_1, b_2, b_3 \in \{0,1\}, \quad (16)$$

$$b_1 + b_2 + b_3 = 1, \quad (17)$$

$$-2 \times b_1 - 3 \times b_2 - 11 \times b_3 \le z \le -b_1 - 2 \times b_2 - 3 \times b_3, \quad (18)$$

$$c_{appr} = (0.09 \times b_1 + 0.009 \times b_2 - 1.25 \times 10^{-4} \times b_3) \times z + (0.19 \times b_1 + 0.028 \times b_2 - 1.375 \times 10^{-3} \times b_3), \quad (19)$$

$$M_{cm} \cdot c_{appr} \le C_{upper}, \quad (20)$$

where $b_1$, $b_2$, and $b_3$ are three $m \times 1$ vectors of instrumental variables, $c_{appr}$ is a $m \times 1$ vector of approximate metabolic concentrations (Fig. S5). Using linear constraints (16) to (20) instead of the non-linear constraints (10) and (14) could substantially increase the computing speed.

Finally, to guarantee that flux configurations we computed can support the survival of a cell, we set a lower bound of the biomass synthesis flux as following:

$$v_{biomass} \ge 1, \quad (21)$$

where $v_{biomass}$ is the flux of biomass synthesis, which is included in all genome-scale metabolic network models.

Overall, TFBA with upper limit of approximate concentration contains the following constraints:

$$S \cdot v = 0, \quad (1)$$

$$v_l \le v \le v_u, \quad (2)$$

$$a \in \{0,1\}, \quad (3)$$

$$(a-1) \times 1000 \le v \le a \times 1000, \quad (4)$$

$$M = 4000, \quad (6)$$

$$-a \times M + \tau \le \Delta_r G \le (1-a) \times M - \tau, \quad (7)$$

$$\tau = 0.05, \quad (8)$$

$$\Delta_r G = \Delta_r G^{\circ} + RTln(10) S^T \cdot z, \quad (9)$$

$$z_l \le z \le z_u, \quad (11)$$

$$\mu_i \le \Delta_r G_i^{\circ} - \Delta_r G_{p,i}^{\circ} \le \upsilon_i, \{i = 1,2,\ldots,n \,|\, \Delta_r G_{p,i}^{\circ} \ is \ known\}, \quad (12)$$

$$\Delta_r G^{\circ} = S^T \cdot \Delta_f G^{\circ}, \quad (13)$$

$$b_1, b_2, b_3 \in \{0,1\}, \quad (16)$$

$$b_1 + b_2 + b_3 = 1, \quad (17)$$

$$-2 \times b_1 - 3 \times b_2 - 11 \times b_3 \le z \le -b_1 - 2 \times b_2 - 3 \times b_3, \quad (18)$$

$$c_{appr} = (0.09 \times b_1 + 0.009 \times b_2 - 1.25 \times 10^{-4} \times b_3) \times z + (0.19 \times b_1 + 0.028 \times b_2 - 1.375 \times 10^{-3} \times b_3), \quad (19)$$

$$M_{cm} \cdot c_{appr} \le C_{upper}, \quad (20)$$

$$v_{biomass} \ge 1, \quad (21)$$

**Predicting directionality of reactions in GEM with TFBA**

All labels of directions of intracellular reactions in the GEM were removed before the application of TFBA to predict their directionalities. Next, we computed the ranges of net flux carried by each reaction by solving the two optimization problems below:

$$\max \text{or} \min \quad v_i\,, \tag{22}$$

where $v_i$ is the flux of the reaction $i$. A positive maximum value means it can happen in the forward direction, and a negative minimum value means it can happen in the backward direction.

Note that the flux direction of a reaction in TFBA framework was determined by two factors, thermodynamics and topology of network. The latter affects whether the metabolites that participate in the reaction could be produced or consumed by other reactions while satisfying all flux balance constraints in the model. Considering the imperfections of the metabolic network, we also computed the range of Gibbs energy change for a reaction by solving another two optimization problems below:

$$\max \text{or} \min \quad \Delta_r G_i\,, \tag{23}$$

where $\Delta_r G_i$ is the Gibbs energy change of reaction $i$. The potential directions of reactions could be inferred by the range of $\Delta G_i$. A positive minimum value of $\Delta_r G_i$ indicates that the reaction can only happen in the backward direction, while a negative maximum value of $\Delta_r G_i$ means that the reaction can only happen in the forward direction.

According to the directionalities inferred based on the range of $v_i$ and $\Delta_r G_i$, we screened all reactions in the GEM, compared the build-in label with the inferred direction, and assigned the direction manually if they were incompatible. The Gurobi[1] solver was used for solving those MIP problems.

**Supplementary Figures**

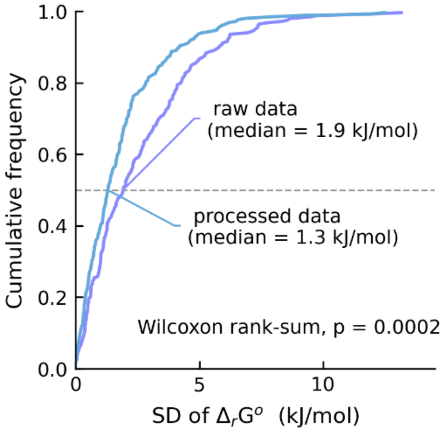


**Fig. S1 (Related to Fig. 1). Distribution of standard deviation of standard Gibbs energy change of reactions.** The mauve line plots the cumulative frequency of raw data and the blue line plots the cumulative frequency of processed data.

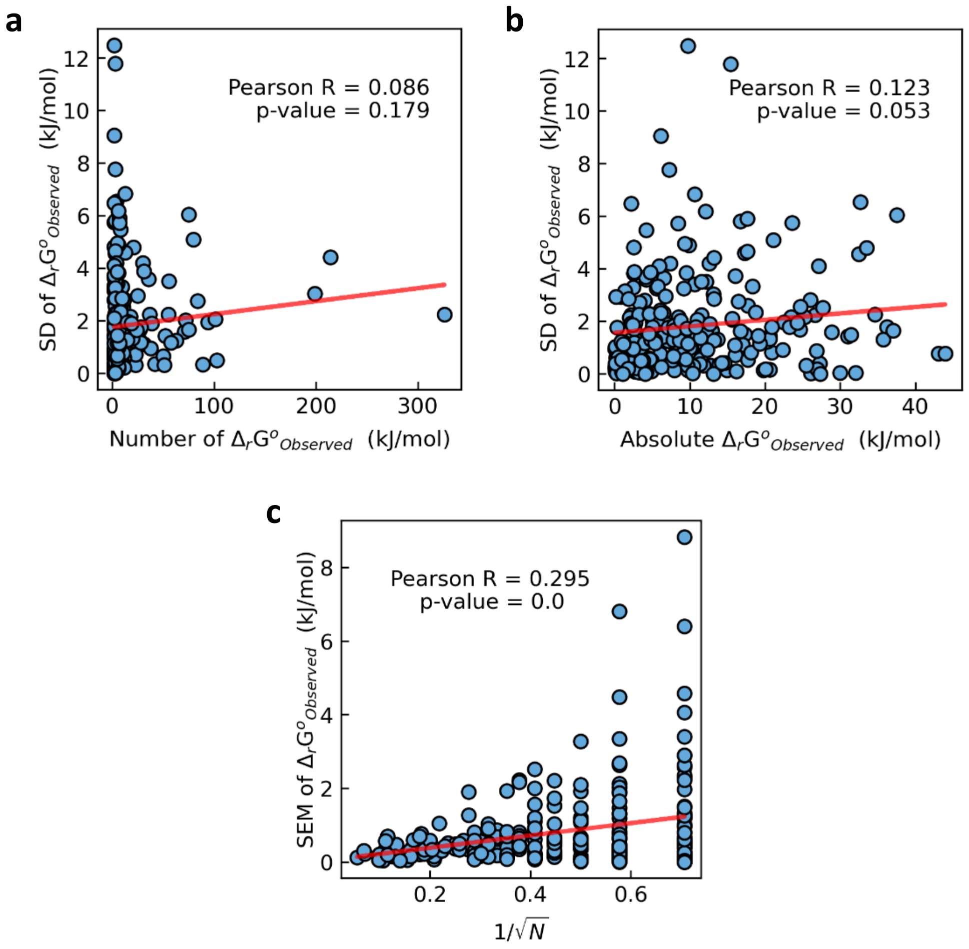


**Fig. S2 (Related to Fig. 1) The relationship between variation of data and other features. a** and **b**, there is no strong correlation between the standard deviation (SD) of reaction's observed Gibbs energy values and other data features. **c**, the significant correlation between the standard error of the mean (SEM) and number of samples suggests that the mean of observed values are convergent when sample sizes increased.

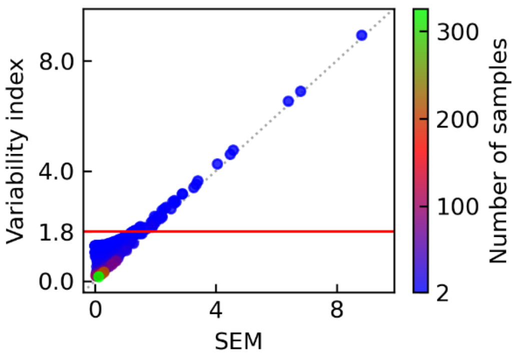


**Fig. S3 (Related to Fig. 1) The relationship between the variability index, SEM, and number of samples.** The variability index of a data item is close to its SEM when the SEM or number of samples is large, otherwise close to the average of standard deviation of total items. Red line indicates the average of standard deviation.

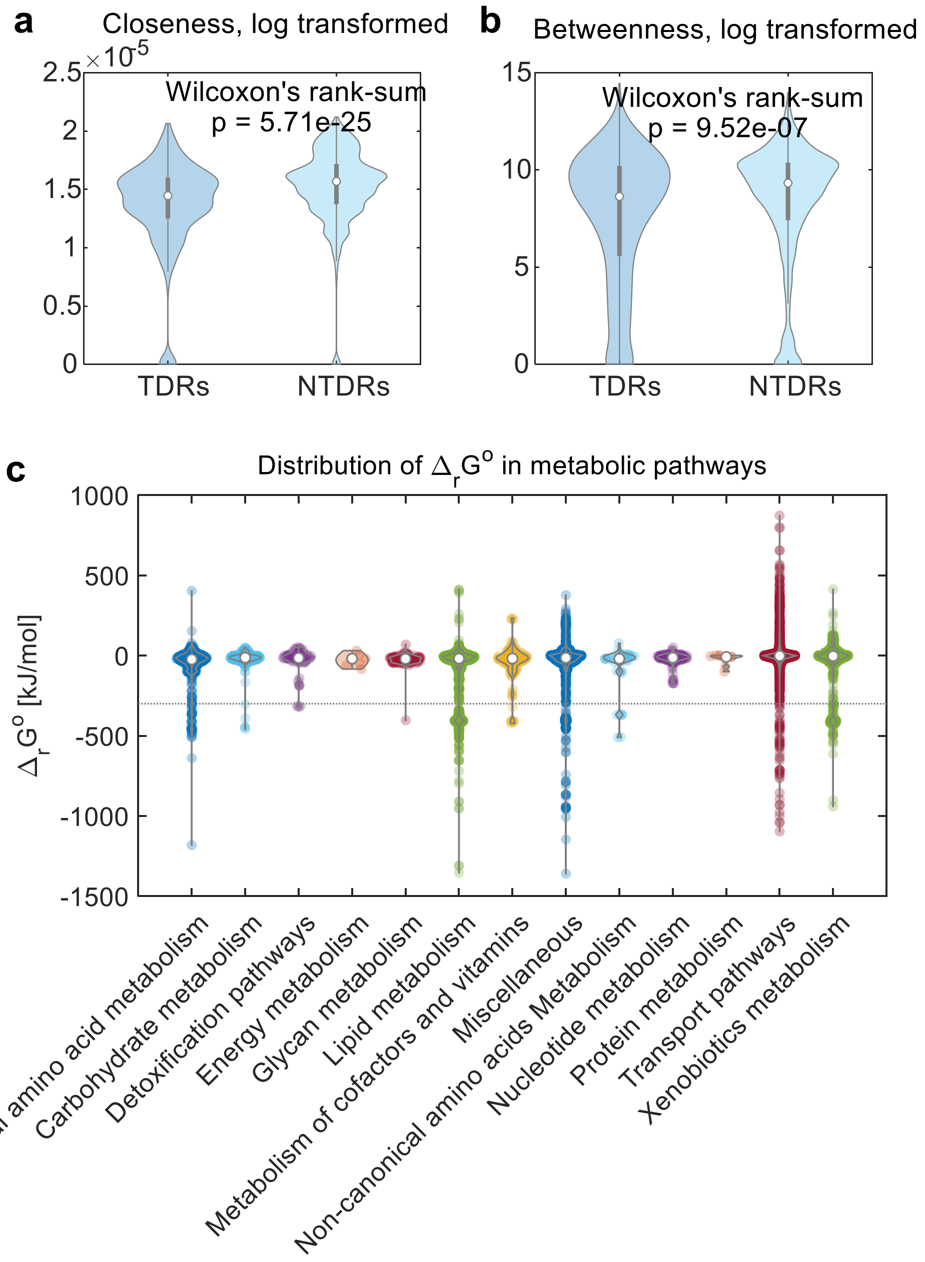


**Fig. S4 (Related to Fig. 5) Distribution of thermodynamic driver reactions (TDRs) in human genome-scale metabolic network.** (a) Violin plots comparing closeness centrality of TDRs and NTDRs. (b) Violin plots comparing betweenness centrality of TDRs and NTDRs. © Violin plots comparing the distributions of standard Gibbs free energy change across metabolic pathways.

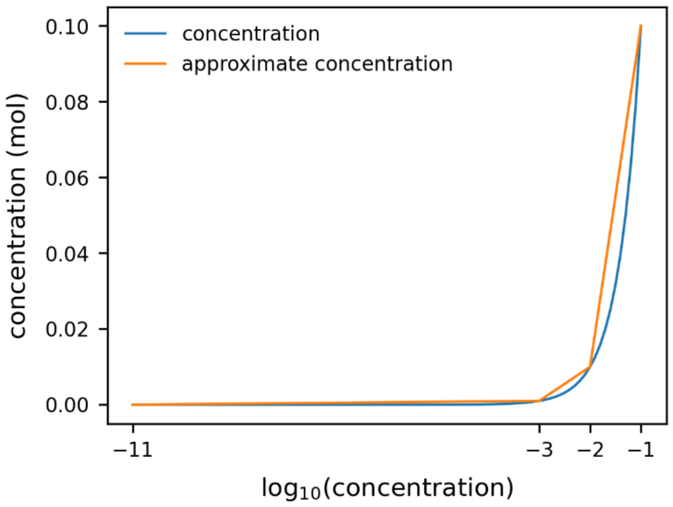


**Fig. S5 (Related to Fig. 6) The actual concentration (blue line) and approximate concentration calculated from piecewise linear approximation of the denary logarithm of concentration (orange line).**

### Supplementary Tables

**Table S1 | Training data statistics (Change of data number in the process of cleaning)**

| | Raw data | High-quality data | Balanced reactions | Unique reactions | Unique metabolites |
|---|---|---|---|---|---|
| TECRDB[2] | 4544 | 3817 | 3814 | 453 | 521 |
| Table of $\Delta_f G^{\circ}$[3] | 223 | 223 | - | - | 222 |

- Not suitable

**Table S2 | List of compound atom and bond features**

| | |
|---|---|
| **Atom features** | |
| Atomic number | 1 ~ 119 |
| Hybridization | unspecified, s, sp, sp2, sp3, sp2d, sp3d, sp3d2, other |
| Aromaticity | not aromatic, aromatic |
| Charge | -4 ~ +4 |
| **Bond feature** | |
| Bond type | unspecified, single, double, triple, quadruple, quintuple, hextuple, one and a half, two and a half, three and a half, four and a half, five and a half, aromatic, ionic, hydrogen, three- center, dativeone, dative, dativel, dativer, other, zero |

One-hot encoding was used for all atom and bond features.

### References

1. Gurobi Optimization, L. (2023).
2. Goldberg, R.N., Tewari, Y.B. & Bhat, T.N. Thermodynamics of enzyme-catalyzed reactions--a database for quantitative biochemistry. *Bioinformatics* **20**, 2874-2877 (2004).
3. Noor, E., Haraldsdottir, H.S., Milo, R. & Fleming, R.M. Consistent estimation of Gibbs energy using component contributions. *PLoS Comput Biol* **9**, e1003098 (2013).
